# Red Palm Oil (RPO) May Enhance Chronic Inflammation Through Disturbance in Lipid Metabolism

**DOI:** 10.1101/2022.09.28.509863

**Authors:** Ridwan Badmos Binyameen, Franklin Akinola

**Affiliations:** Ladoke Akintola University of Technology

## Abstract

Several studies have been published on lipid lowering effect of red palm oil (RPO) with little known about health impact of differential dosage. In this study, we examined lipid profile of hyperlipidemia-induced wistar rats fed with varying quantity of RPO supplementation (15ml/kg, 20ml/kg and 25ml/kg feed). A total of 30 male wistar rats were procured and randomly divided into five groups (A, B, C, D and E) with 6 rats in each group. Group C, D and E received 15 ml/kg, 20ml/kg and 25ml/kg of RPO respectively (thoroughly mixed with high fat diet). No force feeding or oral gavage procedures were employed. While rats in group A were fed with standard rat chow, group B animals fed on high fat diet only and neither of the two groups received RPO supplementation. Plasma concentration of total cholesterol (TC), triglycerides (TG), Albumin, low density lipoproteins (LDL-C), high density lipoproteins (HDL-C), and total proteins (TP) were assessed at the end of the experiment that lasted 4 weeks. In addition to the lipid lowering effect observed in RPO supplementation groups (C, D, E) compared to fatty diet fed only (group B) as widely reported in many studies, both the LDL-C and TG appeared to rise with more RPO supplementation. Findings also revealed the lipid lowering effect more pronounced on triglycerides than the low density lipoproteins. TP in group E was significantly higher compared to group A and B (P < 0.05) and RPO supplementation had a tendency to increase plasma TP.

## INTRODUCTION

The study of dietary fats and oils consumption has gained recognition during the last twenty years, and still continues to especially with emergence of dramatic expansion in areas of metabolic physiology and nutrition science, primarily to combat risks of excess fat related disorders and attendant dysmetabolism. This is because obesity, insulin resistance, and metabolic syndrome X are all linked to diets with high fat content especially the low density lipoprotein (LDL). Various chemicals that make up lipids exhibit a unique characteristic of being soluble in organic solvents but insoluble in water with fatty acids as building block just as amino acids are to protein.

Following this, will it be rational to jettison all fat containing diets for fear of potential harmful effects? Certainly not.

It turns out that fats are not that really bad as presented in many academic gazettes. The truth is, lack of fats in diets is as dangerous as consuming excess of it. Because of this, they are increasingly recognized for their importance and regulatory roles in numerous physiological functions. Lipids and fatty acids act as components of membranes, energy supply and fuel storage, although certain lipids serve other specific purposes. For instance, some act as cell signaling and very low concentrations could cause disturbance in homeostasis resulting in fatal disease conditions like schizophrenia, Alzheimer’s disease, cardiovascular diseases, glomerulonephritis etc. (Bester et al., 2010).

One most important type of lipid that is tightly controlled is the cholesterol. Essentially, and except biological derangement subsists in denovo synthesis, only exogenous cholesterol has proven terribly challenging for the body regulatory mechanisms and learning to regulate daily intake is now a necessity. A number of methods have been identified as interventions, with diet, drugs and exercise of various types and intensities being most favored. However, and for the purpose of this research, we investigated effect of dietary RPO supplementation on lipid profile in hyperlipidemia-induced model of wistar rats.

Excerpts from people’s feeding habits reveal attitude as major challenge in related diet-controlled experiments in human subjects. Unlike animals, human beings are hard to put and maintain under experimental conditions. Many would not like to forsake preferred diets regardless of disease burden that might ensue. Even when recruited for experiments of this nature, they hardly abide to prescribed nutrition regimen or daily calorie modification. This may be one of the reasons many studies continue to report conflicting findings with so much discrepancy that make it practically impossible to appreciate the true impacts of varieties of diet interventions. To control for this, we resorted to animal study and investigated one of the most widely consumed cooking oil ‘red palm oil’ having reviewed substantial literatures on its lipid lowering potential.

RPO is extracted from a tropical plant Elaeis guineensis (commonly known as the oil palm) which produces second largest volume of vegetable oil in the world (Eidangbe et al., 2010). Overall biochemical effects of RPO’s most important group constituents which are fatty acids and antioxidants like carotenoids, tocopherol and tocotrienols have been well debated with a view to harness possible therapeutic values in treatment of obesity and hyperlipidemia conditions. Notwithstanding, presence of high percentage of saturated fatty acids (up to 51%) has been cited against its lipid lowering potential and reduction in cardiovascular risk factors (CVR) in obese patients. Hence, we hypothesize that palm oil does not affect lipid profile in diet-induced hyperlipidemia in wistar rats model.

## RESULTS

### EXPERIMENTAL DESIGN

Thirty male wistar rats were procured from the Anatomy Department, Faculty of Basic Medical Sciences, Ladoke Akintola University of Technology, Ogbomoso, Oyo State Nigeria. Rats were weighed and initial weights recorded. After two weeks of acclimatization period, they were further randomly divided and assigned into five groups with six rats in each group. Animals were housed in plastic cages specially designed for this experiment with roof made of net for adequate ventilation. They were daily fed to satisfaction and had free access to water and food throughout four weeks the experiment lasted.

| Groups | Treatment (water ad libitum for all groups) |
| --- | --- |
| A | Standard rat chow for 4 weeks (negative control). |
| B | Fatty diet only for 4 weeks (positive control). |
| C | Fatty diet + RPO (15ml/kg feed) for 4 weeks. |
| D | Fatty diet + RPO (20ml/kg feed) for 4 weeks. |
| E | Fatty diet + RPO (25ml/kg feed) for 4 weeks. |

Rats in groups B, C, D and E were fed high fat diet to induce hyperlipidemia while those in group A were fed standard rat chow. In addition, group C, D, and E were fed with fatty diets supplemented with locally refined RPO purchased at Sabo market, Ogbomoso, Oyo State. The RPO was thoroughly mixed with experimental diets to avoid texture concern. No force feeding or oral gavage. Body weight of the animals were recorded on weekly basis until end of the experiment.

### NUTRITIONAL CONTENT OF THE HIGH FAT EXPERIMENTAL DIET

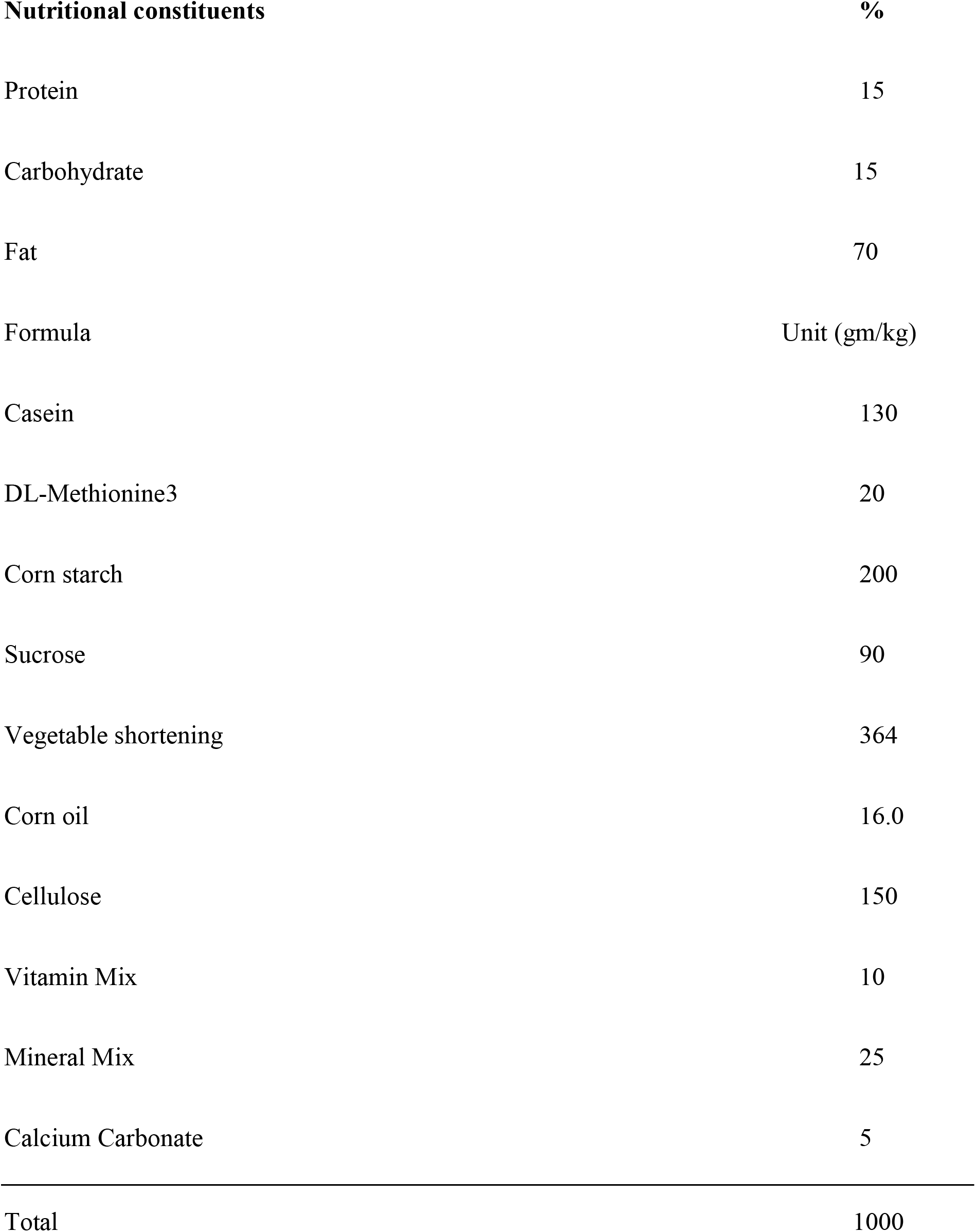

| **Nutritional constituents** | **%** |
| --- | --- |
| Protein | 15 |
| Carbohydrate | 15 |
| Fat | 70 |

| **Formula** | **Unit (gm/kg)** |
| --- | --- |
| Casein | 130 |
| DL-Methionine<sup>3</sup> | 20 |
| Corn starch | 200 |
| Sucrose | 90 |
| Vegetable shortening | 364 |
| Corn oil | 16.0 |
| Cellulose | 150 |
| Vitamin Mix | 10 |
| Mineral Mix | 25 |
| Calcium Carbonate | 5 |
| Total | 1000 |

### BLOOD COLLECTION AND STORAGE

Blood samples were collected from the heart by terminal bleeding. Venous blood was withdrawn from each rat at the end of the study and collected into sample bottles containing heparin anticoagulant. Samples were centrifuged and analyzed at the Bioethics Laboratory, Institute for Advanced Medical Research and Training (IMRAT), University College Hospital (UCH) Ibadan, Oyo State, Nigeria.

### PLASMA PREPARATION

Blood collected into heparin sample bottles were gently mixed and centrifuged at 4000g for 10mins and supernatants collected into plain bottles for analysis.

### METHODOLOGY

#### ESTIMATION OF PLASMA TRIGLYCERIDES

Plasma triglycerides was determined using GPO-PAP colorimetric method according to Jacobs et al., (1960).

#### ESTIMATION OF PLASMA TOTAL CHOLESTEROL

Total cholesterol was estimated using the enzymatic Endpoint method according to Allain et al., (1974)

#### ESTIMATION OF PLASMA HDL AND LDL

Estimation of plasma HDL and LDL was done using CHOD-PAP method according to Bairaktari et al., (2000). In summary, low density lipoproteins (LDL and VLDL) and chylomicron fraction were precipitated quantitatively by the addition of phosphotungstic acid in the presence of magnesium ions. After centrifugation, the cholesterol concentration in the HDL fraction, which remains in the supernatant, was determined using Friedewald formula (Friedewald, 1972).

#### ESTIMATION OF PLASMA TOTAL PROTEIN (TP)

Total protein was estimated using Biuret Method (colorimetric end point) according to Lubran, (1978).

#### ESTIMATION OF PLASMA ALBUMIN

Plasma albumin was determined by the method of Doumas and Watson, (1981). Briefly, measurement was based on its quantitative binding to the indicator 3, 3’, 5, 5’-tetrabromocresol sulphonepphthalein (bromocresol green, BCG). The albumin-BCG-complex absorbs maximally at 578nm, the absorbance being directly proportional to the concentration of albumin in the sample.

### STATISTICAL ANALYSIS

SPSS package was used for all statistical analyses. Data were expressed as mean± standard error of mean(SEM). The statistical analysis was carried out using (ANOVA) and *Schiff’s post hoc and* level of significance was accepted at (p<0.05).

## TABLES

**Table 1:**
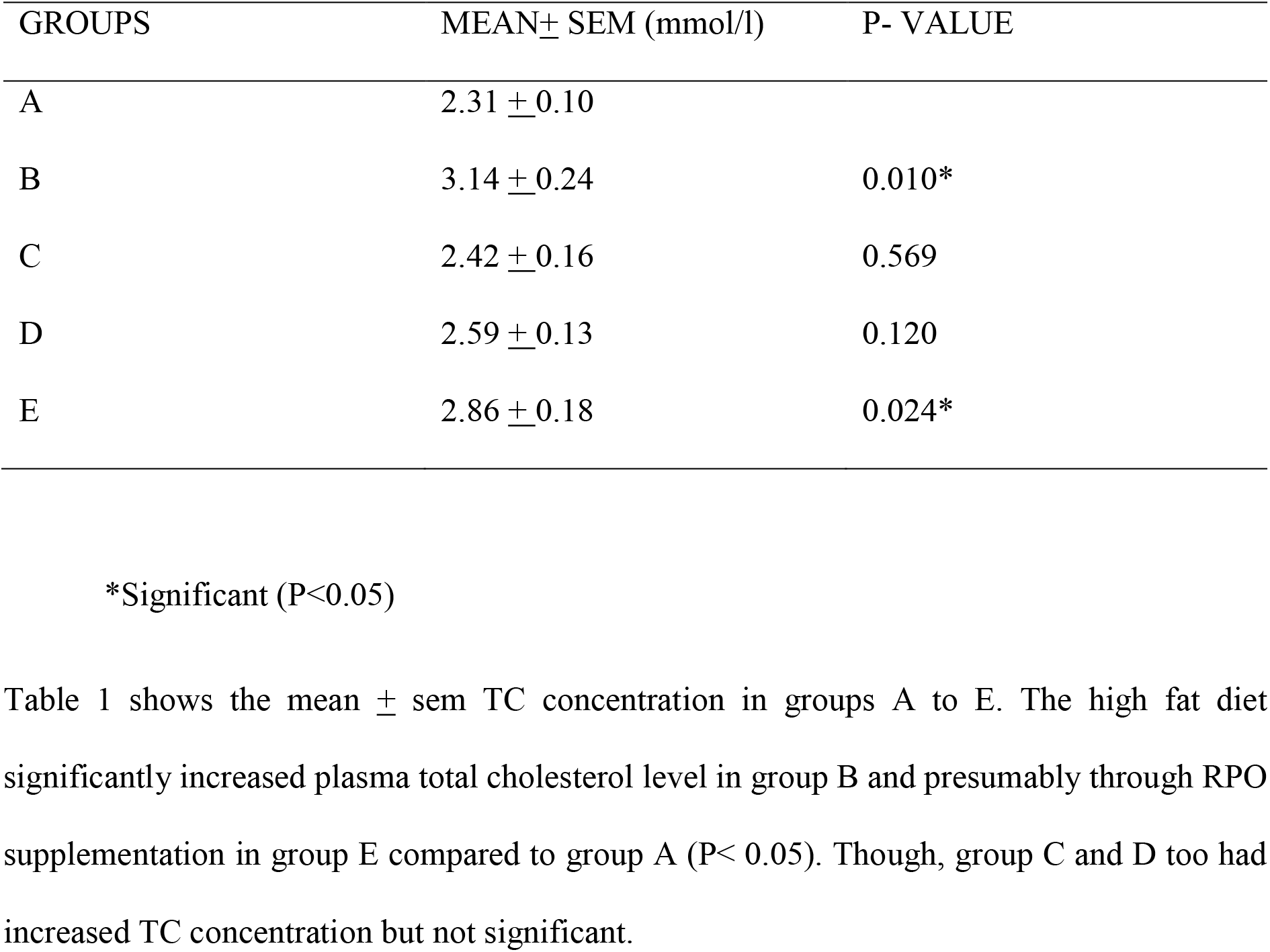
MEAN ± SEM Total Cholesterol (TC) comparison Between Group A and other Groups.

**TABLE 2:** MEAN ± SEM Total Cholesterol(TC) comparison between group B and C, D, E.

| GROUPS | MEAN± SEM (mmol/l) | P- VALUE |
| --- | --- | --- |
| B | 3.14 ± 0.24 | |
| C | 2.42 ± 0.16 | 0.033* |
| D | 2.59 ± 0.13 | 0.074 |
| E | 2.86 ± 0.18 | 0.374 |

Table 2 shows the mean ± sem TC concentration. Plasma TC level was significantly lower in group C compared to other experimental groups ( $P < 0.05$ ) that received RPO.

**TABLE 3:** MEAN ± SEM Total Cholesterol (TC) comparison between group C and D, E.

| GROUPS | MEAN± SEM (mmol/l) | P- VALUE |
| --- | --- | --- |
| C | 2.42 ± 0.16 | |
| D | 2.59 ± 0.13 | 0.439 |
| E | 2.86 ± 0.18 | 0.103 |

Table 3 shows the mean ± sem plasma TC concentration of group C compared to D and E. There was no statistical difference between TC level among compared groups.

**TABLE 4:** MEAN ± SEM Total Cholesterol comparison between group D and E.

| GROUPS | MEAN± SEM (mmol/l) | P- VALUE |
| --- | --- | --- |
| D | 2.59 ± 0.13 | |
| E | 2.86 ± 0.18 | 0.261 |

Table 4 presents the mean ± sem plasma TC concentration of group D compared with E. There was no statistical difference in TC level between the two groups.

**Table 5:** MEAN ± SEM Total Protein (TP) comparison between group A and other groups.

| GROUPS | MEAN± SEM (g/dl) | P- VALUE |
| --- | --- | --- |
| A | 54.00 ± 2.59 | |
| B | 55.00 ± 2.42 | 0.681 |
| C | 53.00 ± 3.31 | 0.877 |
| D | 65.00 ± 1.87 | 0.006* |
| E | 65.00±1.77 | 0.005* |

Table 5 presents the mean ± sem plasma TP concentration. A difference was observed in mean plasma TP concentration across the group ( $P < 0.05$ ). However, a statistically significant increase in TP level was only observed in groups D and E compared to A ( $P < 0.05$ ).

**Table 6:** MEAN ± SEM Total Protein (TP) comparison between group B and C, D, E.

| GROUPS | MEAN± SEM (g/dl) | P- VALUE |
| --- | --- | --- |
| B | 55.00 ± 2.42 | |
| C | 53.00 ± 3.31 | 0.609 |
| D | 65.00 ± 1.87 | 0.010* |
| E | 65.00±1.77 | 0.008* |

Table 6 presents the mean ± sem of TP concentration. Plasma TP was significantly higher in group D and E compared to group B ( $P < 0.05$ ).

**Table 7:** MEAN ± SEM Total Protein (TP) comparison between group C and D, E.

| GROUPS | MEAN± SEM (g/dl) | P- VALUE |
| --- | --- | --- |
| C | 53.00 ± 3.31 | |
| D | 65.00 ± 1.87 | 0.011* |
| E | 65.00± 1.77 | 0.010* |

Table 7 shows the mean ± sem plasma total protein concentration. Group D and E have significantly higher plasma TP compared to group C ( $P < 0.05$ ).

**Table 8:** MEAN ± SEM Total Protein(TP) comparison between group D and E.

| GROUPS | MEAN± SEM (g/dl) | P- VALUE |
| --- | --- | --- |
| D | 65.00 ± 1.87 | |
| E | 65.00± 1.77 | 0.950 |

Table 8 presents comparison between group D and E. No significant difference was observed in plasma TP between the groups

**Table 9:** MEAN ± SEM Triglycerides (TG) comparison between group A and other groups.

| GROUPS | MEAN± SEM (mmol/l) | P- VALUE |
| --- | --- | --- |
| A | 0.41 ± 0.16 | |
| B | 0.96 ± 0.18 | 0.045* |
| C | 0.64 ± 0.11 | 0.272 |
| D | 0.69 ± 0.09 | 0.162 |
| E | 0.73±0.14 | 0.163 |

**Table 10:** MEAN ± SEM Triglycerides (TG) comparison of group B with C, D and E.

| GROUPS | MEAN± SEM (mmol/L) | P- VALUE |
| --- | --- | --- |
| B | 0.96 ± 0.18 | |
| C | 0.64 ± 0.11 | 0.162 |
| D | 0.69 ± 0.09 | 0.210 |
| E | 0.73±0.14 | 0.333 |

Table 10 shows the mean ± sem plasma TG. There was no significant difference in TG level observed between the groups.

**Table 11:** MEAN ± SEM Triglycerides (TG) comparison of group C with group D and E.

| GROUPS | MEAN± SEM (mmol/L) | P- VALUE |
| --- | --- | --- |
| C | 0.64 ± 0.11 | |
| D | 0.69 ± 0.09 | 0.740 |
| E | 0.73±0.14 | 0.626 |

Table 11 shows the mean ± sem of plasma TG concentration of group C compared with group D and E. No statistical difference is seen in TG level among the groups.

**Table 12:** MEAN ± SEM Triglycerides (TG) comparison between group D and E.

| GROUPS | MEAN± SEM (mmol/L) | P- VALUE |
| --- | --- | --- |
| D | 0.69 ± 0.09 | |
| E | 0.73±0.14 | 0.815 |

Table 12 shows the mean ± sem plasma TG concentration of group D compared with group E. Group E has higher plasma TG though not significant.

**Table 13:** MEAN ± SEM Plasma Albumin comparison between group A and other groups.

| GROUPS | MEAN± SEM (g/dl) | P- VALUE |
| --- | --- | --- |
| A | 26.17 ± 1.82 | |
| B | 27.00 ± 2.57 | 0.796 |
| C | 27.00 ± 2.44 | 0.789 |
| D | 30.83 ± 1.49 | 0.074 |
| E | 32.67± 1.43 | 0.018* |

**Table 14:**
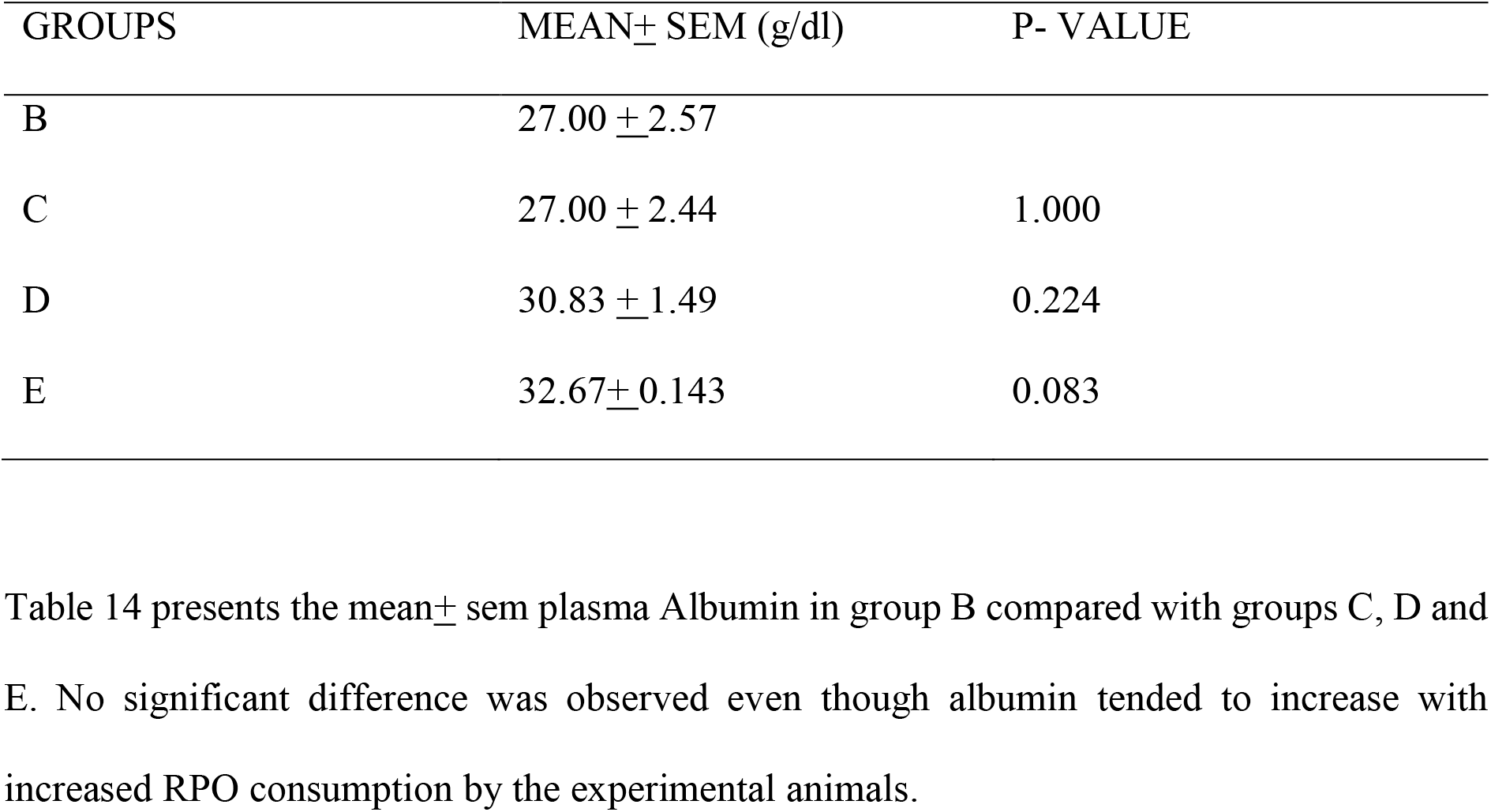
MEAN ± SEM Plasma Albumin comparison between group B and group C, D, E.

| GROUPS | MEAN± SEM (g/dl) | P- VALUE |
| --- | --- | --- |
| B | 27.00 ± 2.57 | |
| C | 27.00 ± 2.44 | 1.000 |
| D | 30.83 ± 1.49 | 0.224 |
| E | 32.67± 0.143 | 0.083 |

Table 14 presents the mean ± sem plasma Albumin in group B compared with groups C, D and E. No significant difference was observed even though albumin tended to increase with increased RPO consumption by the experimental animals.

**Table 15:** MEAN ± SEM Plasma Albumin comparison of group C and group D, E.

| GROUPS | MEAN± SEM (g/dl) | P- VALUE |
| --- | --- | --- |
| C | 27.00 ± 2.44 | |
| D | 30.83 ± 1.49 | 0.288 |
| E | 32.67± 0.143 | 0.073 |

**Table 16:** MEAN ± SEM Plasma Albumin comparison between group D and E.

| GROUPS | MEAN± SEM (g/dl) | P- VALUE |
| --- | --- | --- |
| D | 30.83 ± 1.49 | |
| E | 32.69± 0.143 | 0.392 |

**Table 17:** MEAN ± SEM HDL-C comparison between group A and other groups.

| GROUPS | MEAN± SEM (mmol/dl) | P- VALUE |
| --- | --- | --- |
| A | 1.42 ± 0.07 | |
| B | 1.24 ± 0.15 | 0.296 |
| C | 1.33 ± 0.16 | 0.635 |
| D | 1.43 ± 0.10 | 0.924 |
| E | 1.15± 0.21 | 0.257 |

Table 17 shows the mean ± sem HDL-C between the groups A-E. No significant difference in plasma HDL-C in group A compared with B-E.

**Table 18:** MEAN ± SEM HDL-C comparison between group B and group C, D, E.

| GROUPS | MEAN± SEM (mmol/dl) | P- VALUE |
| --- | --- | --- |
| B | 1.24 ± 0.15 | |
| C | 1.33 ± 0.16 | 0.675 |
| D | 1.43 ± 0.10 | 0.304 |
| E | 1.15±0.21 | 0.738 |

Table 18 shows the mean ± sem HDL-C. No significant difference observed in plasma HDL-C when group B was compared with C, D, E.

**Table 19.** MEAN ± SEM HDL-C comparison between group C and D, E.

| GROUPS | MEAN± SEM(mmol/dl) | P-VALUE |
| --- | --- | --- |
| C | 1.33±0.16 | |
| D | 1.43± 0.10 | 0.615 |
| E | 1.15± 0.21 | 0.507 |

Table 19 shows the mean ± sem HDL-C. No significant difference observed in HDL-C in group C when compared with D and group E.

**Table 20:**
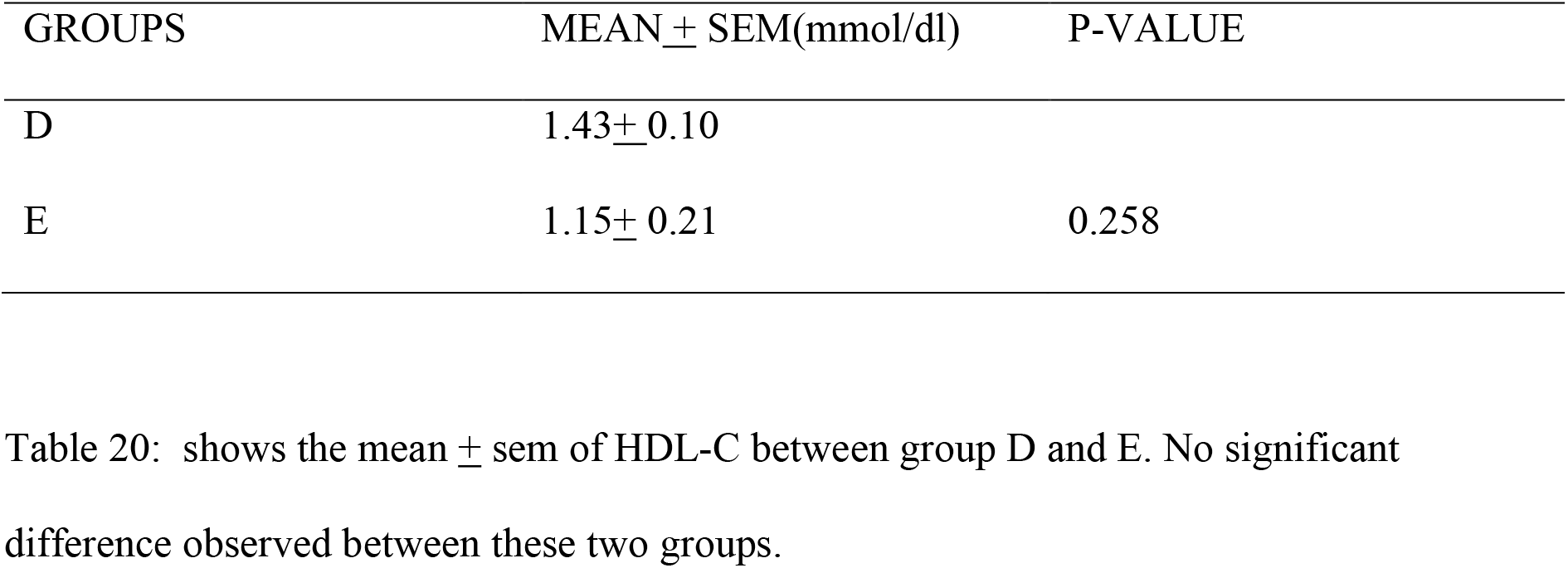
MEAN ± SEM HDL-C comparison between group D and E.

**Table 21:**
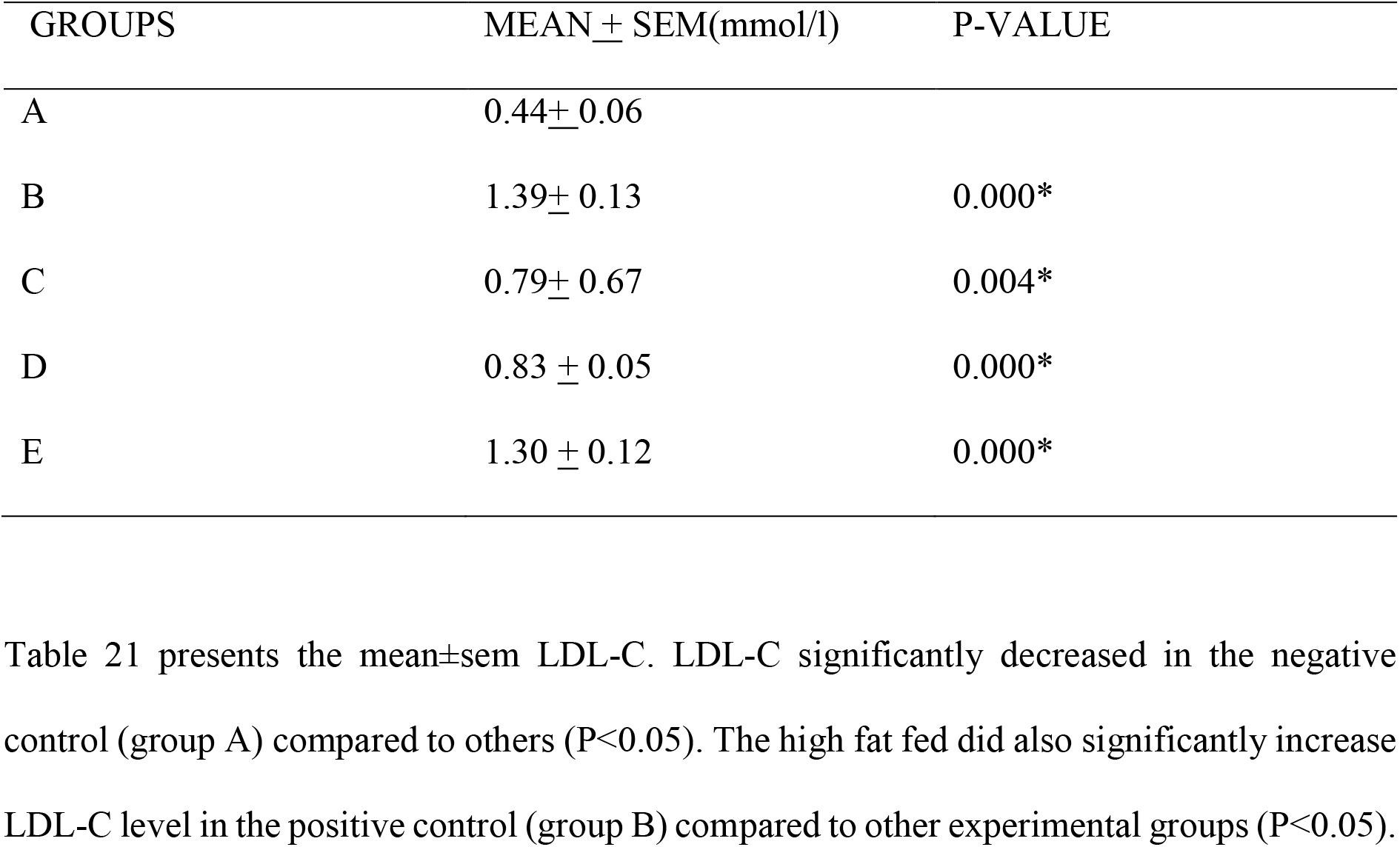
MEAN ±SEM LDL-C comparison of Group A with other groups.

**Table 22:** MEAN ±SEM LDL-C comparison of group B with C, D and E.

| GROUPS | MEAN ± SEM(mmol/l) | P-VALUE |
| --- | --- | --- |
| B | 1.39± 0.13 | |
| C | 0.79± 0.07 | 0.002* |
| D | 0.83± 0.05 | 0.003* |
| E | 1.30 ± 0.12 | 0.643 |

**Table 23:** MEAN + SEM LDL-C comparison of group C with D and E.

| GROUPS | MEAN ± SEM(mmol/l) | P-VALUE |
| --- | --- | --- |
| C | 0.79± 0.07 | |
| D | 0.83± 0.05 | 0.594 |
| E | 1.03 ± 0.12 | 0.003* |

Table 23 shows the mean ± sem of LDL-C. Significant difference was noted in group C when compared with group E ( $P < 0.05$ ). However, no significant difference between group C that received lowest dose of RPO (15ml/kg fatty diet) and group D that received higher dose of 20ml of RPO per kg fatty diet. Note that, significant difference occurred only in group E that received highest dose (25ml of palm oil/ kg fatty diet).

**Table 24:** MEAN ± SEM LDL-C comparison between group D and E.

| GROUPS | MEAN± SEM (mmol/l) | P- VALUE |
| --- | --- | --- |
| D | 0.83 ± 0.05 | |
| E | 1.30±0.12 | 0.004* |

**TABLE 25:** MEAN ± SEM weight gain between group A-E showing effects of the high fat diet and palm oil supplementation.

| WEIGHT(gm) | GROUP A | GROUP B | GROUP C | GROUP D | GROUP E |
| --- | --- | --- | --- | --- | --- |
| INITIAL | 47.33±0.62 | 47.33±0.62 | 61.83±0.79 | 66.83±0.70 | 65.50±1.23 |
| FINAL | 90.33±1.82 | 164.50±6.24 | 124.83±3.43 | 123.83±3.88 | 132.67±2.38 |
| WEIGHT<br>GAIN | 43±1.92 | 117.17±6.43 | 63±3.52 | 57±3.94 | 67.17±2.68 |

**Table 26:** MEAN ±SEM weight gain comparison of Group A with other groups.

| GROUPS | MEAN ± SEM(g) | P-VALUE |
| --- | --- | --- |
| A | 43±1.92 | |
| B | 117.17± 6.43 | 0.000* |
| C | 63±3.52 | 0.001* |
| D | 57± 3.94 | 0.01* |
| E | 67.17 ± 2.68 | 0.000* |

The above table compared the mean weight gain in grams of group A to others. Group A, B, C, D and E had significantly higher mean weight ( $p < 0.05$ ).

**Table 27:** MEAN ±SEM weight gain comparison of Group B with C, D and E.

| GROUPS | MEAN ± SEM (g) | P-VALUE |
| --- | --- | --- |
| B | 117.17±6.43 | |
| C | 63±3.52 | 0.000* |
| D | 57±3.94 | 0.000* |
| E | 67.17 ± 2.68 | 0.000* |

The above table compared the mean weight gain in grams of group B to C, D and E. Group B animals have raised mean value for weight gain that is significantly different from C, D and E.

**Table 28:** MEAN ±SEM weight gain comparison of group C with D and E.

| GROUPS | MEAN± SEM(g) | P-VALUE |
| --- | --- | --- |
| C | 63±3.52 | |
| D | 57±3.94 | 0.283 |
| E | 67.17± 2.68 | 0.368 |

The above table presents information about weight gain in group C compared to D and E. Group D that received 20ml/kg RPO supplementation had lower mean weight change compared to E (25ml/kg RPO supplementation) and C (15ml/kg RPO supplementation).

**Table 28:**
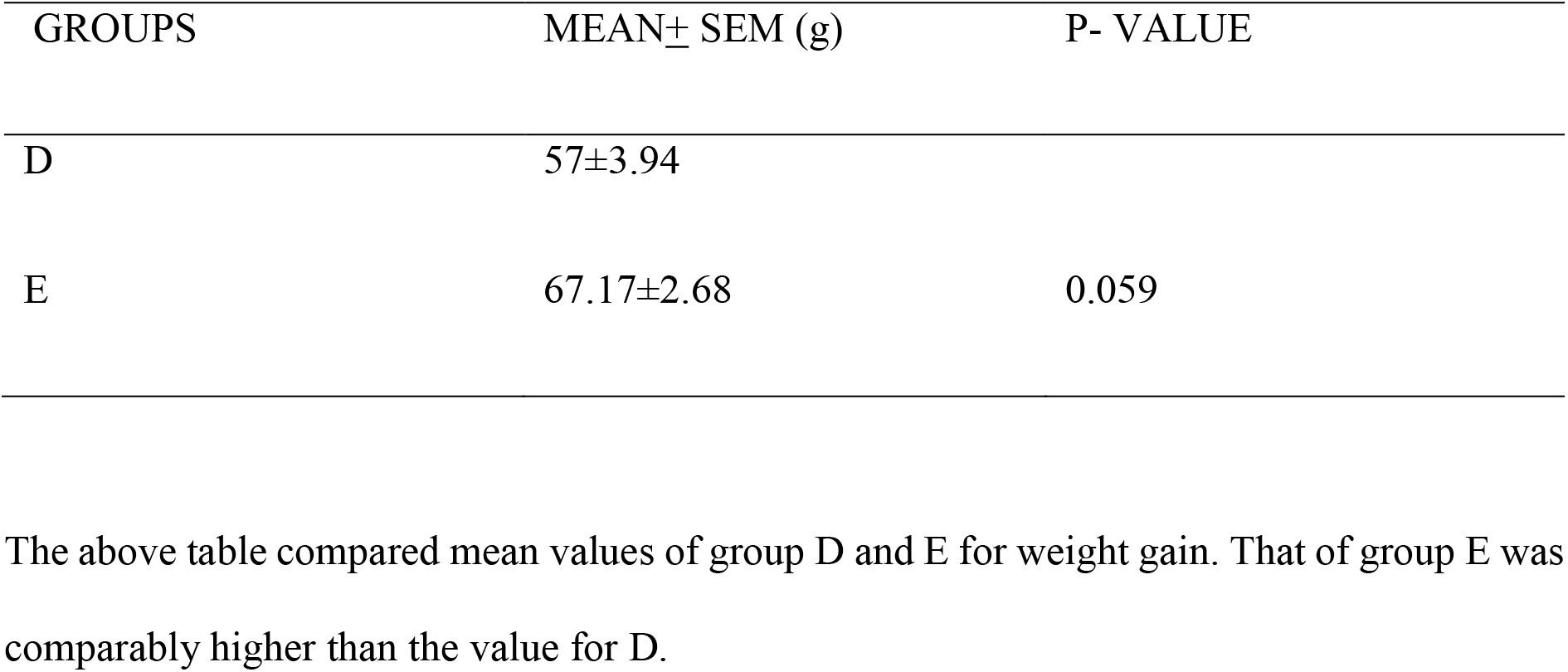
MEAN ±SEM weight gain comparison of group D with E.

| GROUPS | MEAN± SEM (g) | P- VALUE |
| --- | --- | --- |
| D | 57±3.94 | |
| E | 67.17±2.68 | 0.059 |

The above table compared mean values of group D and E for weight gain. That of group E was comparably higher than the value for D.

**Figure :1.**
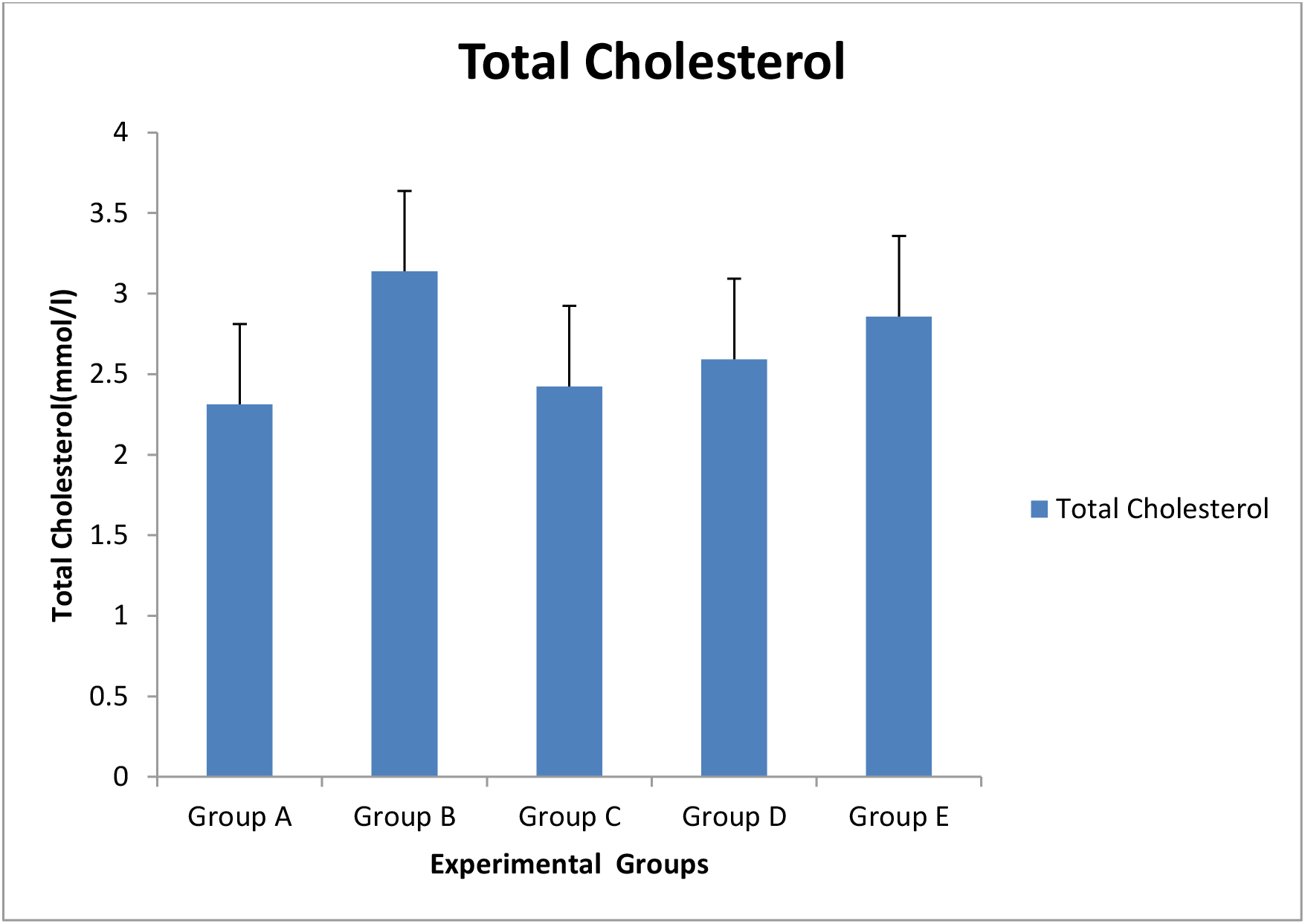
Graph showing comparison of mean plasma TC among experimental groups

**Figure: 2.**
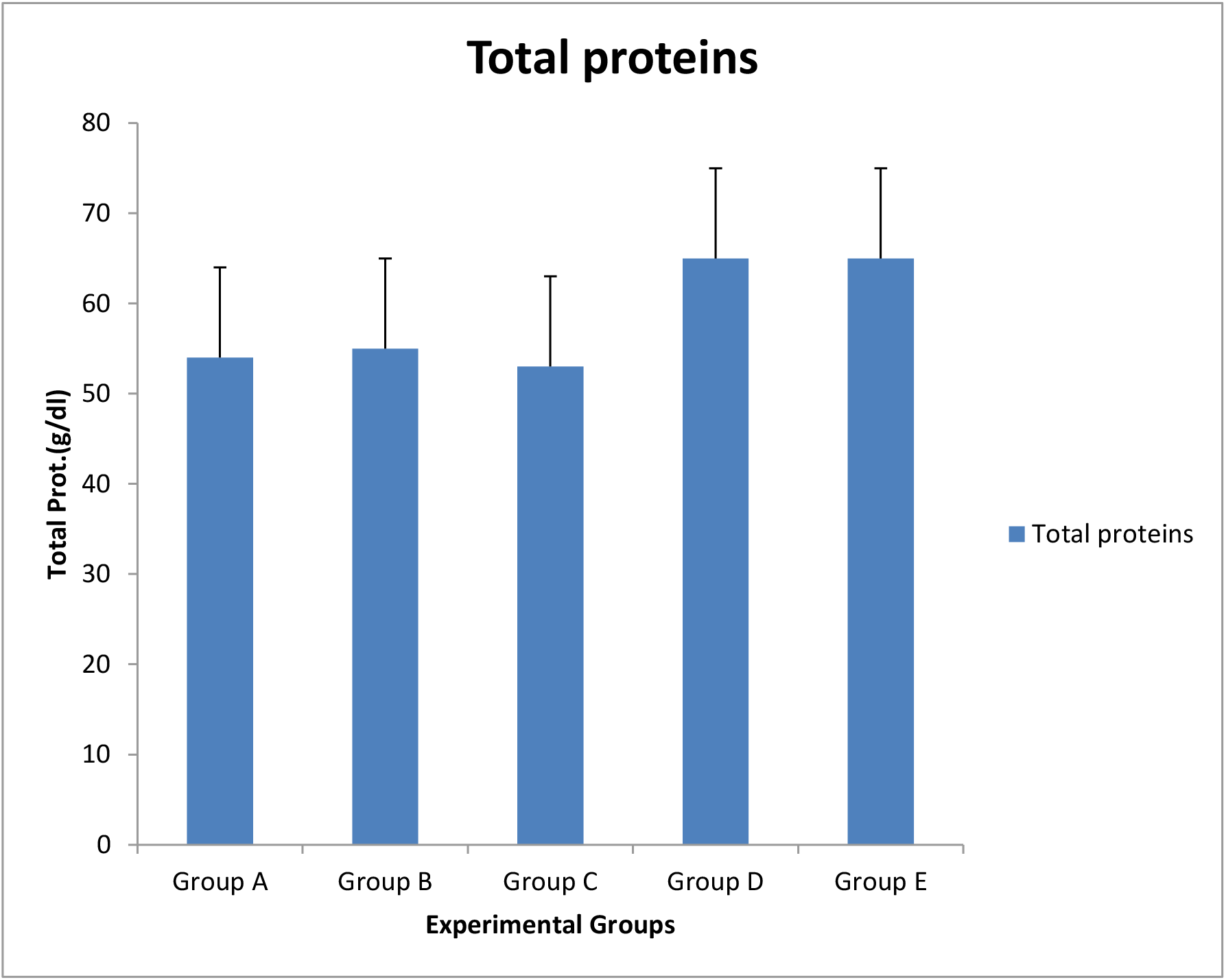
Graph showing comparison of mean plasma TP among experimental groups

**Figure: 3.**
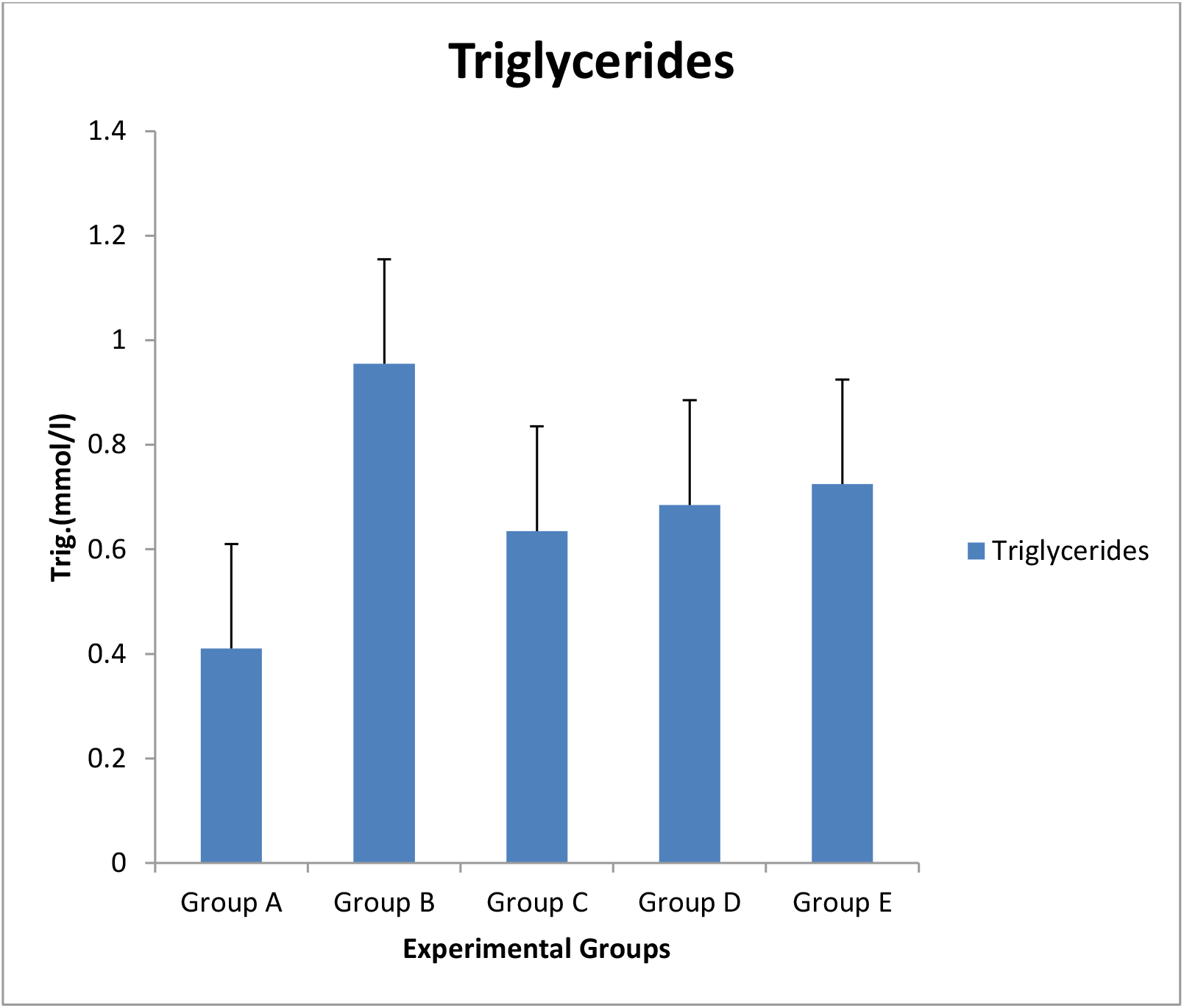
Graph showing comparison of mean plasma TG among experimental groups

**Figure: 4.**
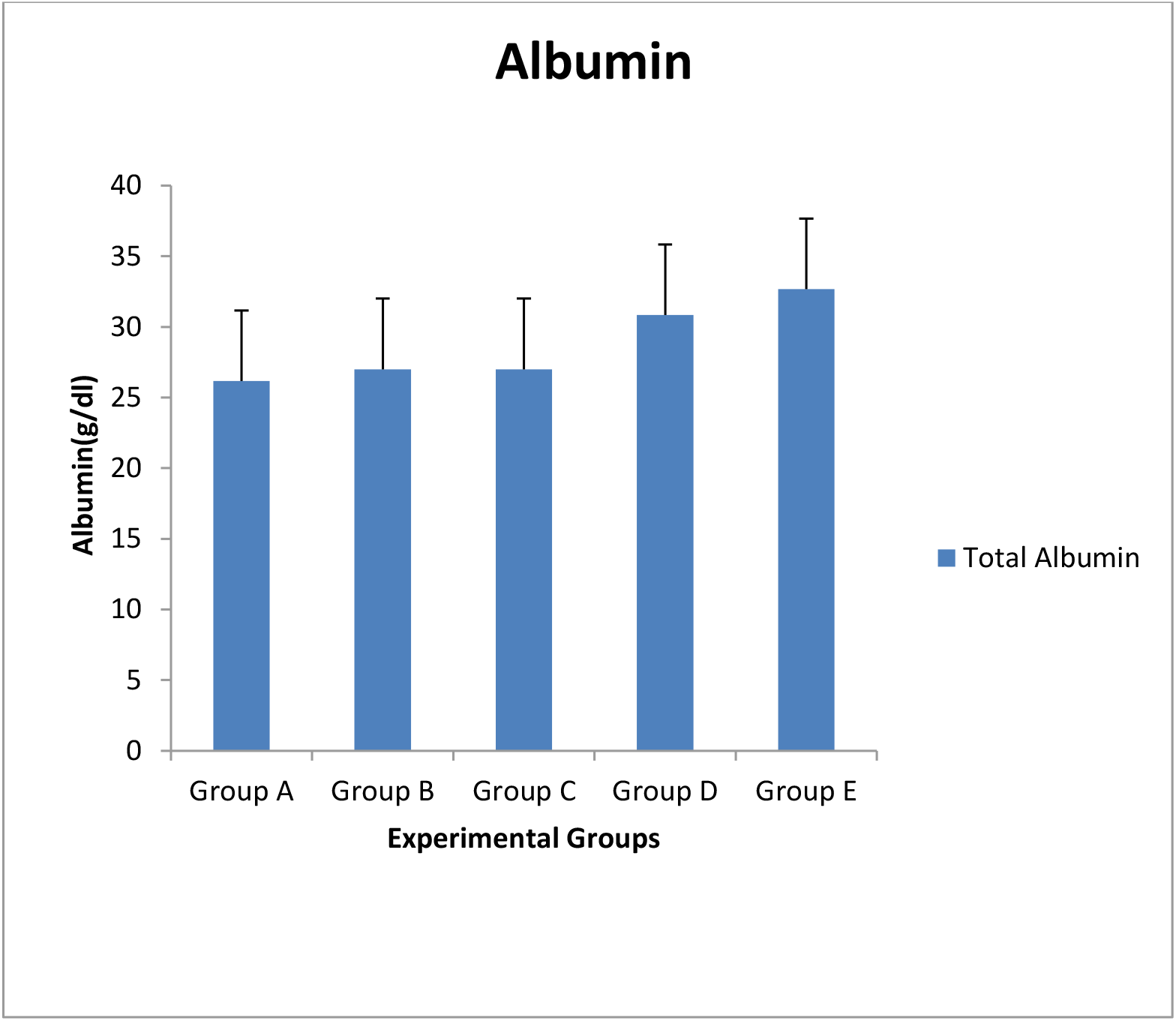
Graph showing comparison of mean plasma albumin among experimental groups

**Figure: 5.**
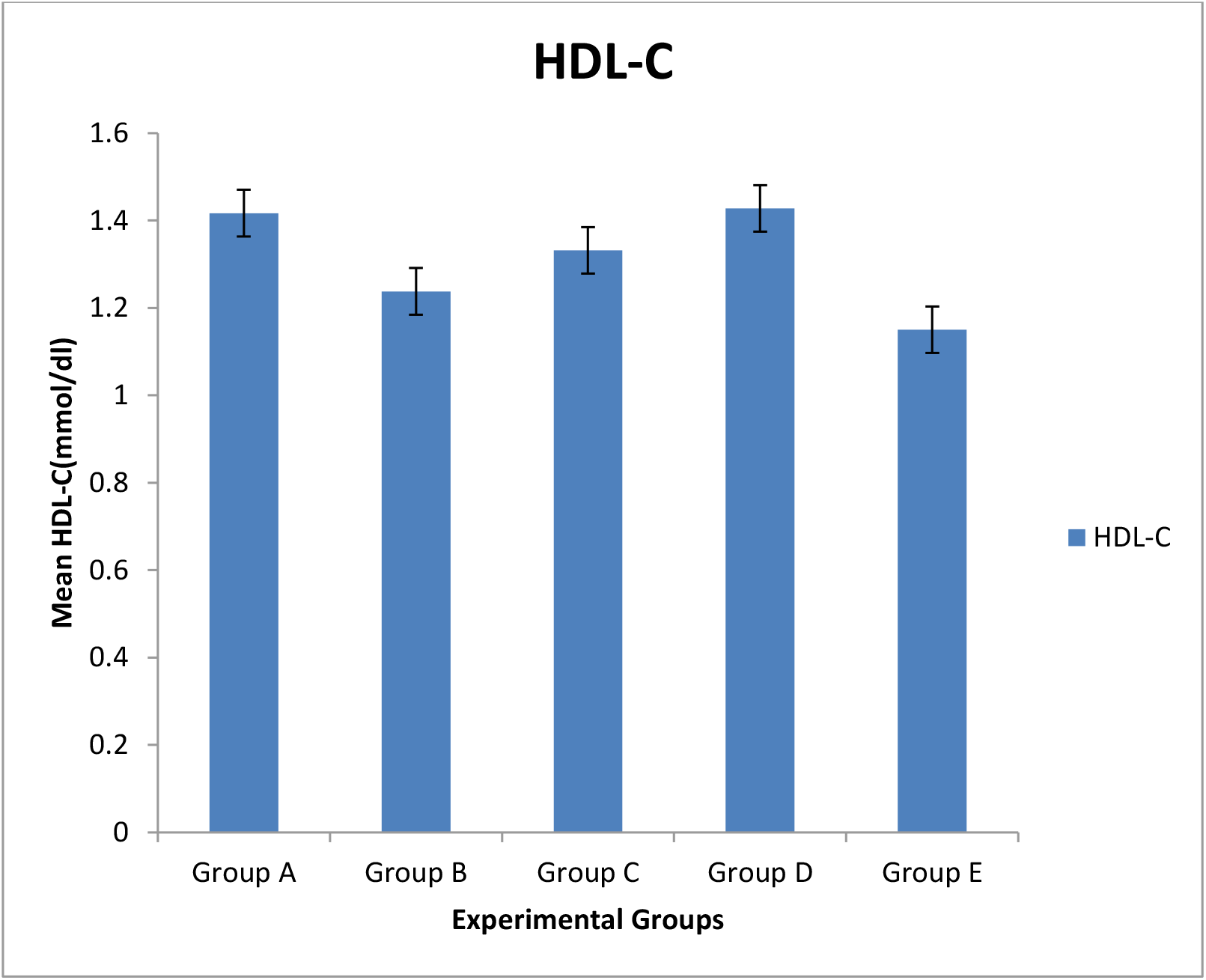
Graph showing comparison of mean plasma HDL-C among experimental groups

**Figure: 6.**
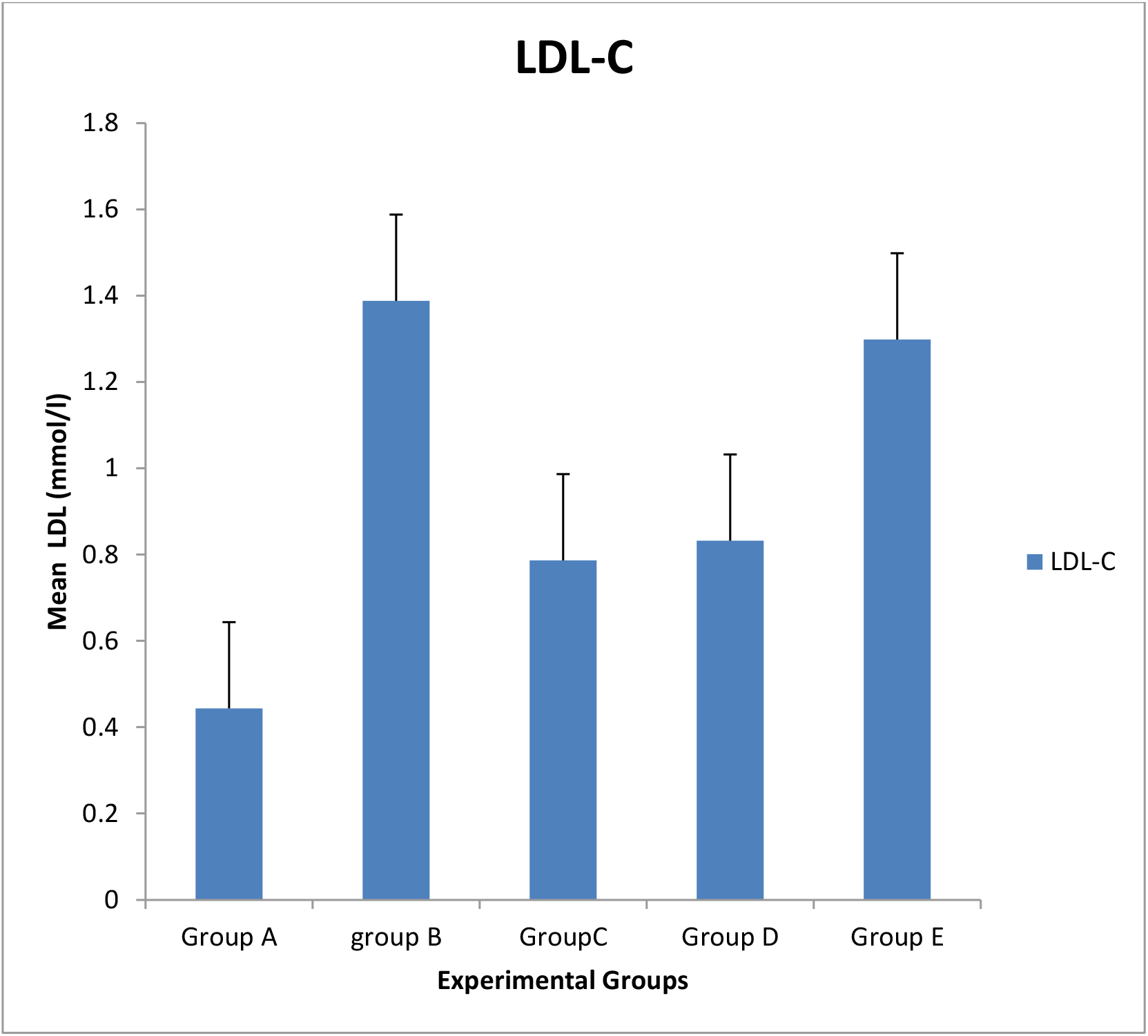
Graph showing comparison of mean plasma LDL-C among experimental groups

**Figure: 7.**
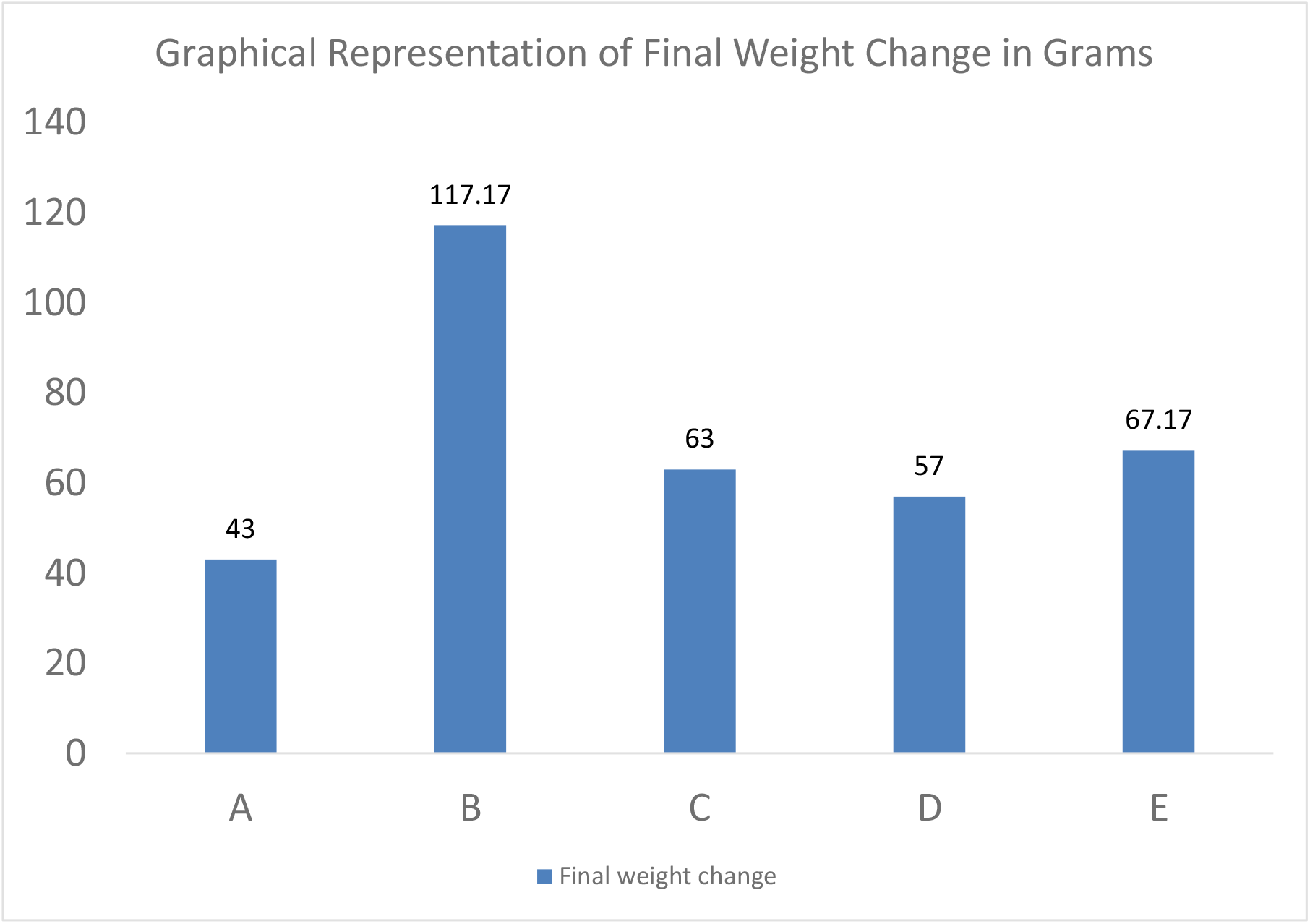
Graph showing comparison of mean weight gain among experimental groups

**Figure: 8.**
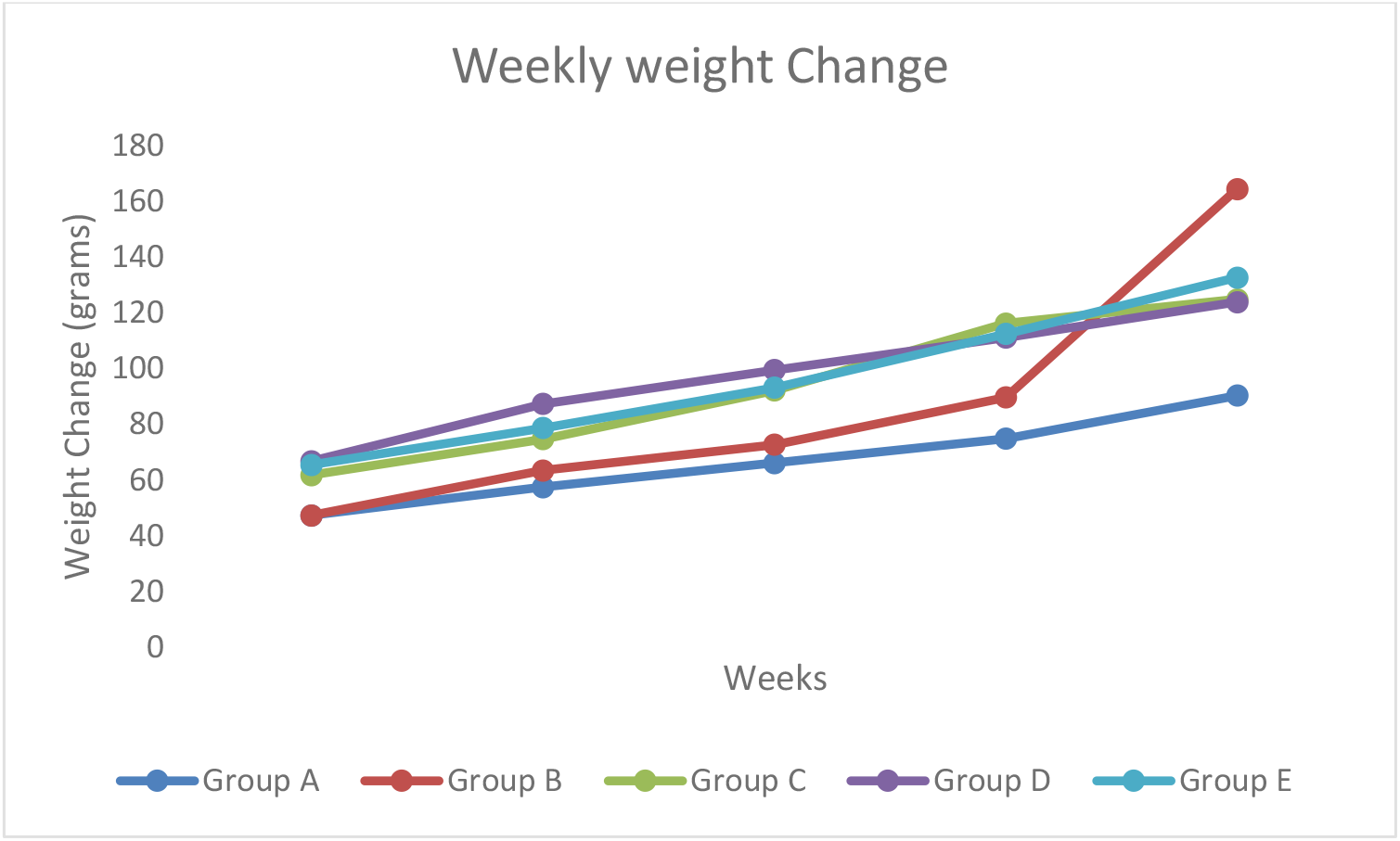
Graph showing weekly change in mean weight among experimental groups

## DISCUSSION AND CONCLUSION

### DISCUSSION

In this study, we found gradual but concomitant increase in plasma TG and LDL-C as RPO supplementation increases. The mechanism by which RPO aids lipid accumulation remains unresolved among authors investigating health impacts. But diets rich in saturated fatty acids (SFA) like RPO are deemed more obesogenic compared to ones with less saturated or polyunsaturated fatty acids (PUFA) (Hariri and Thibault, 2010) and higher serum lipid with RPO was reported by Choi et al., (2014). Although, long-term study shows plasma level remains fairly constant (Manorama and Rukmini, 1991). Significantly higher weight gain observed in fatty diet fed group without RPO fortification compared to groups with RPO supplementation confirms lipid lowering effect of RPO as reported by Oguntibeju et al., (2009), Eidangbe et al., (2010), Ayeleso et al., (2012) and Syarifah-Noratiqah et al., (2020) albeit believed to primarily center on prevention of rise in TG and LDL-C level. But tendency of RPO to boost lipid profile at a later time in a dose-dependent manner (as noted in the present study) has received less attention and remains a major public health concern.

Interestingly, sustained high plasma LDL-C and TG triggers cardiovascular disease risks, having been implicated in a wide range of dysmetabolism as evidenced in several studies. Thus, we are not surprised that plasma albumin follows similar trend as a major transporter for the abundant free fatty acids (FFA). Less soluble in aqueous solution of the body fluid, short to medium chain fatty acids are regularly transported by albumin whose concentration indirectly determines plasma lipid availability (Vusse, 2009; Alharthi et al., 2019).

In normal condition of homeostasis, the highly reactive cholesterol is shielded from longtime interaction with body tissues. Intracellular lipid droplets, denovo synthesis, uptake by circulating lipoproteins, storage and extracellular excretion are held under tight control by providing safe passage through esterification. The body employs the esterification process to store and transport cholesterol (both dietary and synthetic) effectively while as well protecting cells against toxicity of excess (otherwise free or unesterified) cholesterol (Gonen and Miler, 2020). Although, lipoproteins in this process, are capable of absorbing minute free cholesterol restricted to their outer surface. More cholesterol can, however, be packed inside of them (safe corridor) as inert cholesteryl esters which significantly expands lipoproteins capacity and makes it possible for cholesterol to easily transverse the bloodstream. But this mechanism is most effective for lipid regulation during homeostasis and in conjunction with the HDL-C.

The HDL-C is functionally antithetical to LDL-C by acting to continuously adjust and restore the critical biological equilibrium making its plasma concentration an important health determining factor. Unfortunately, its restoration effect is often attenuated when overwhelmed by external factors that favor formation of more LDL-C than it is able to offload as apparent in the present study. This further proves RPO may lack the high-end therapeutic values attributed to it in many studies as HDL-C did not improve with further supplementation.

During and prior to metabolic derangement, however, LDL-C (tagged bad cholesterol) is aggressively mobilized to tissues from areas like the digestive system and liver even when they seem waring of it. The very low density lipoprotein (VLDL) is, in a similar manner, notorious for enriching tissues with both LDL-C and TG. It primarily effects this by packing TG and cholesteryl esters (CEs) together then later shed the former after undergoing steady transformational remodeling. The creeping pile-up in plasma TG and LDL-C observed in this study with RPO increment suggests high plasma VLDL and possible malicious health burden (such as intramuscular lipid infiltration) capable of progressing into full blown age-dependent sarcopenic obesity characterized by impaired muscle power, strength and performance (Poggigiogalle et al., 2022). And by implications, since abundance of and feedback from lipids and metabolites upregulate pyruvate dehydrogenase kinase-4 (PDK4) and up tissue dependence on them for energy and utilization (Tuika et al., 2012; Crewe et al., 2013; Zhang et al., 2014), elevated plasma lipid has been described as one material precursor for oxidative stress and type2 diabetes as less glucose are metabolized leading to insulin resistance (Lichtenstein and Schwab, 1999; In-Kyu Lee 2013), cardiovascular diseases (Crewe al., 2013) and arrays of other metabolic dysfunctions that follow the initial subclinical inflammation (Nacsimento et Al., 2008).

In addition to albumin that helps transport free fatty acids (FFA), observed tendency of TP to rise with dietary RPO signals low-grade but progressive and potentially damaging oxidative stress in the experimental animals, presumptively through the inflammation sensitive proteins (ISP) like fibrinogen, orosomucoid, alpha-1 antitrypsin, heptoglobin, and ceruloplasmin associated with future weight gain (Engstrom et al., 2003). In the same vein, the ISP may be responsible for the rise in cellular oxidative stress and inflammation reported on repeatedly heated RPO. Along with cholesterol, they have also been linked to myocardial infarction, cardiovascular death, and stroke (Engstrom et al., 2015) suggesting that chronic inflammation may be associated with rise in plasma lipids.

Overall, we found high quality evidence that RPO may initiate, aid and augment subclinical inflammation, hyperlipidemia, atherosclerosis, cardiovascular disease risks, type2 diabetes and stroke in wistar rat model and advise that caution be applied in dietary consumption and daily intake.

#### Concluding Remarks

RPO processing takes place in stages which include threshing, (removal of fruit from bunches), sterilization of bunches, digestion of the fruits, pressing (oil extraction), and finally is clarification and drying. Unfortunately, many local RPO processing facilities as common in Nigeria lack financial capacity to acquire technology that can preserve and protect the unsaponified contents (tocopherols, tocotrienols, carotenes, phytosterols etc.) often praised for lipid lowering effect (Watkins et al., 1993). Hence, the tendency to raise cellular oxidative stress, blood pressure (Leong et al., 2009), TC and cause atherosclerosis (Adam et al., 2008) through excessive heating and overriding effect of the SFA contents. Even though we did not measure plasma antioxidants in this research, we acknowledge its lipid lowering effect and possible agent for differential scientific reportage about lipid lowering potential of RPO.

Furthermore, unprocessed fruit bunches can be kept for days if not weeks in the local processing factories and RPO can be hoarded for months by local marketers to be sold later at expensive price. These factors, among others that we do not have control over, are capable of compromising RPO quality by increasing the level of free fatty acids (FFA) and peroxides through the activities of microbial lipase (Choon-Hui et al., 2009). But for lack of quality testing and control in Nigeria, no one has kept an eye on FFA content of publicly sold RPO talk more of rancidity status. Although, RPO source employed in this experiment was adjudged to be of highest nutritional quality in the Southwestern Nigeria (Akinola et al., 2010). We are aware that local processors often change approach based on economic reality especially when no one watches over them. By the way, the wear and tear of the rusty cooking irons used in the heating process coupled with the locally fabricated milling machines can adulterate final products by raising heavy metal contamination known to compromise the body’s antioxidant defense system (Ichipi-Ifukor1 et al, 2022).

For this, results from RPO studies may vary significantly based on many factors including processing culture, region, technological sophistication, and waste disposal strategy. We stress on importance of differential technological advances on RPO quality which authors have ignored. We propose that future research should consider Malaysian RPO as gold standard against locally refined products in other parts of the world. This will not only help unravel the growing inconsistent findings regarding its therapeutic benefits but also signals policy makers on the product quality consumed by the multitudes and thus assist in strengthening public health systems. The RPO adopted in the study of Ayeleso et al., (2012) that reported decreased lipid profile with increased RPO supplementation differs in source and most probably in nutritional composition from the one employed in the present experiment. Generally, we believe the more advanced refining technology is, the better and healthier is the RPO quality.

## Notes

### Competing Interest Statement

The authors have declared no competing interest.

